# Proteomic analysis of chemosensory organs in the honey bee parasite *Varroa destructor*: a comprehensive examination of the potential carriers for semiochemicals

**DOI:** 10.1101/260539

**Authors:** Immacolata Iovinella, Alison McAfee, Guido Mastrobuoni, Stefan Kempa, Leonard J. Foster, Paolo Pelosi, Francesca Romana Dani

## Abstract

The mite *Varroa destructor* is the major parasite of the honey bee and is responsible for great economical losses. The biochemical tools used by *Varroa* to detect semiochemicals produced by the host are still largely unknown. We have performed proteomic analysis on chemosensory organs of this species in order to identify putative soluble carriers for pheromones and other olfactory cues emitted by the host. In particular, we have analysed forelegs, mouthparts (palps, chelicera and hypostome) and the second pair of legs (as control tissue) in reproductive and phoretic stages of the *Varroa* life cycle. We identified 958 *Varroa* proteins, most of them common to organs and stages. Sequence analysis shows that four proteins can be assigned to the odorant-binding protein (OBP)-like class, which bear some similarity to insect OBPs, but so far are only reported in some Chelicerata. In addition, we have detected the presence of two proteins belonging to the Niemann-Pick family, type C2 (NPC2), which have been suggested to act as semiochemical carriers. This work contributes to elucidating the chemical communication systems in *Varroa* with the aim of understanding how detection of semiochemicals has evolved in terrestrial non-hexapod Arthropoda. Data are available via ProteomeXchange with identifier PXD008679.

## Introduction

One of the main threats to honey bee colonies^1^ worldwide is the mite *Varroa destructor* (hereon referred to as ‘*Varroa*’). Females of this ectoparasite are transmitted between hives by foraging bees, and once in the hive they settle in the bee larval cells and lay eggs. The newborn *Varroa*, generally one male and four females for each cell, feed on the honey bee larvae and, once the females leave the cell, spread in the hive by adhering to adult bees.

Communication between *Varroa* individuals as well as their interactions with honey bees are mediated by chemical signals. Some cuticular hydrocarbons of bee larvae as well as 2-hydroxyhexanoic acid, a component of brood food, have been reported as attractants for mites in their reproductive stage ^2-5^. Once inside the cells, a blend of three fatty acid methyl esters produced by the bee pupae regulates laying of unfertilized (male) and fertilized (female) eggs^6^ by *Varroa*, and induce the reproductive maturation of young *Varroa*^7^. Mature female mites attract males with a cocktail of three fatty acids (palmitic, stearic, and oleic) and their ethyl esters^8^. While in their phoretic stage, the *Varroa* are repelled by geraniol and nerolic acid^9^, as well as by (Z)-8-heptadecene^10^, which are all produced by the foragers; for this reason, the mites tend to parasitize nurse bees.

Compared to insects, chemical communication in other arthropods, particularly Chelicerata, is poorly understood. Most of the studies are focused on morphology^11^ and electrophysiology ^12-14^ while several papers report on the identification of putative semiochemicals^8,9,15-18^. Gustation and olfaction take place in sensilla, which are located on mouthparts and forelegs in ticks and mites. In *Varroa* the main olfactory organ, referred to as pit organ, is located on forelegs and presents nine olfactory hairs, which are morphologically similar to insect sensilla basiconica^19,20^. Furthermore, electrophysiological experiments have clearly demonstrated that the forelegs of *Varroa* respond to chemical stimuli^21,22^.

Only preliminary information is available on *Varroa*’s biochemical tools (receptors and carrier proteins) for chemosensing. Based on genome and transcriptome projects, ionotropic receptors and gustatory receptors have been identified in some ticks and mites^23-26^, but chelicerates lack homologs of the typical insect olfactory receptor family^27,28^. Odorant-binding proteins (OBPs), which act as carriers of odorants and pheromones in the sensillar lymph of insects, are absent in Chelicerata^27^.

The presence of CSPs also seems questionable. A single sequence reported in the tick *I. scapularis*^25^ turned out to be identical with a CSP of the mosquito *Culex quinquefasciatus* (acc. XP_001844693), indicating a result of contamination. Furthermore, the two CSPs reported in a transcriptome study of the mite *Tyrophagus putrescentiae*^29^ are very similar to CSPs of Diptera (around 80% identity), leaving the possibility of contamination an open question. Therefore, in the absence of OBPs and CSPs, other carrier proteins are likely to be present in the chemosensing systems of Chelicerata.

A third family of proteins possibly acting as semiochemical carriers in insects include the NPC2 (Niemann-Pick proteins of type C2) proteins^30-32^. This family is well represented in Chelicerata with a variable number of genes^31,33^, depending on the species. In particular, in the tick *Ixodes scapularis*, a dozen genes have been identified and one of the encoded proteins was detected by immunocytochemistry experiments in chemosensilla of this species^34^. Members of the NPC2 family have been also found in the tick *Amblyomma americanum*^35^ and eight transcripts encoding such proteins have recently been reported in a transcriptome project in *Varroa* chemosensory organs^26^. For NPC2 proteins, a function of semiochemical carriers seems to be well supported by their ligand-binding properties as well as by their localization in chemosensilla^30,32,34^. Moreover, three-dimensional structures of NPC2 members both from vertebrates and insects are available, some of them containing hydrophobic ligands inside their binding pockets^30,36^.

Another class of soluble proteins has been proposed as semiochemical carriers in the tick *A. americanum*^35^ and in two spider species^24^, as well as in *Varroa*^26^. Given some structural similarity with insect OBPs, these proteins have been named as ‘OBP-like.” Sequence identity values with insect OBPs are generally low (around 15% or less) and the pattern of six cysteines, a typical signature of most insect OBPs, is not fully conserved. Some OBP-like proteins of Chelicerata contain four cysteines in a pattern resembling that of insect C-minus OBPs, but other members present six cysteines, although in positions different from those of classic OBPs of insects^35^. Binding data and cellular localization are still needed to support their putative role in chemosensing.

In this work we report the results of a proteomic analysis on chemosensory organs of *Varroa* to better understand chemical communication in this economically devastating species. In particular, knowledge of the molecular mechanisms used by the mites to follow chemical signals from the larval bees could provide the basis for alternative strategies to control the population of the parasite inside the hive.

## Experimental Procedures

### Sample collection

Adult mites were collected at two different stages: 'reproductive mites’ from drone larvae and 'phoretic mites’ from young adult bees, foragers, or adult drones. Specimens were kept at -20°C until dissection. Reproductive mites were collected from frames containing exclusively drone brood, produced by workers after excluding the queen from that part of the frame, in an apiary located in Certaldo (Firenze). Phoretic mites were collected from adult bees in the experimental apiary at the Department of Biology, University of Firenze. Foragers and drones were collected with a net in front of the hive, while young bees were obtained from brood frames temporarily removed from the hive.

Dissections were performed on ice and three appendages were isolated: forelegs, bearing the tarsal organ; mouthparts, containing palps, chelicera and hypostome; and the second pair of legs, to be used as control (Figure 1).

**Figure 1.**
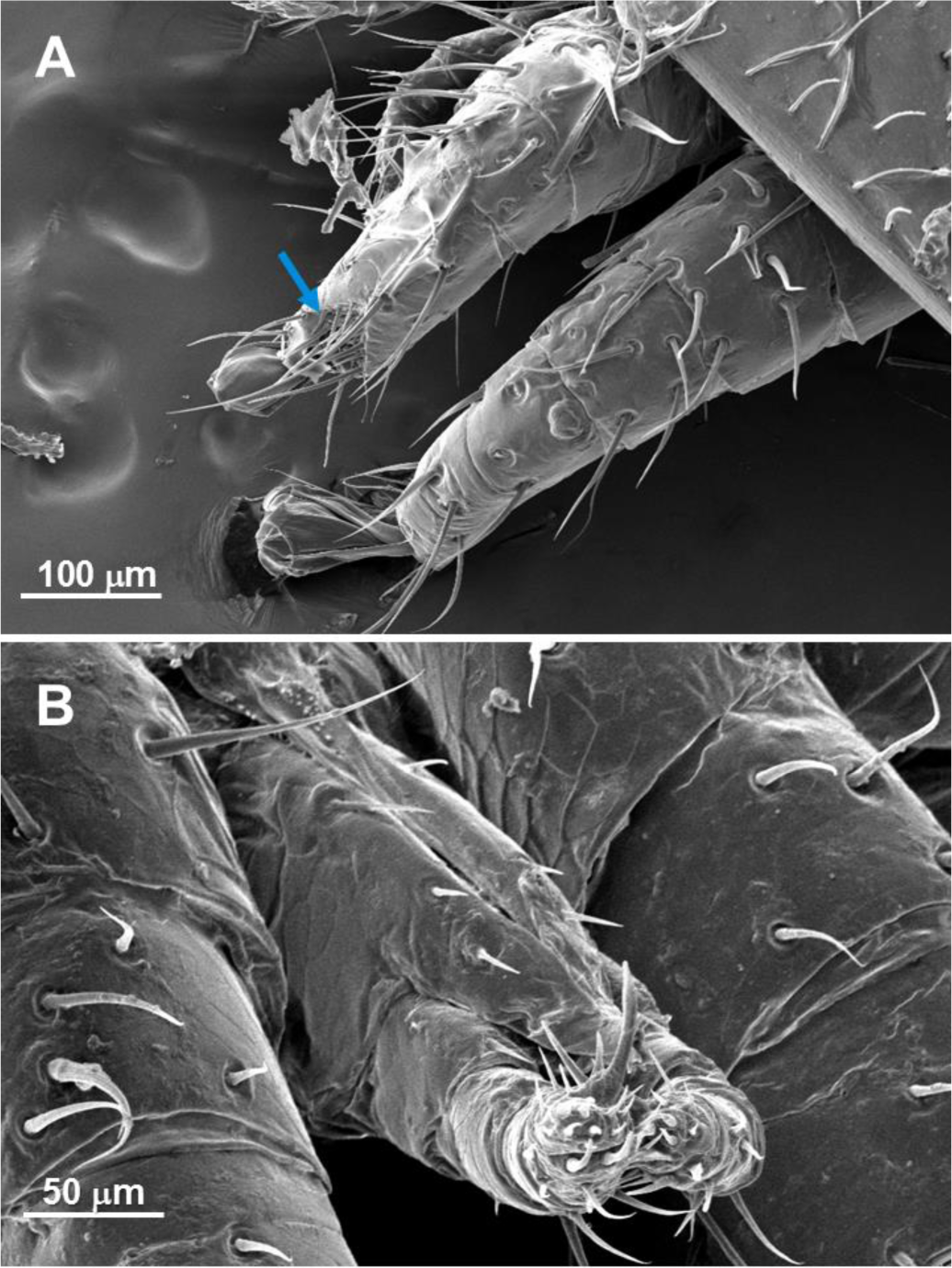
Scanning electron microscope images of forelegs and second pair of legs of an adult female coated with gold (panel A) and ventral view of mouth parts of an adult female coated with graphite and gold (panel B). The chemosensory pit organ is visible on the foreleg tarsi (red arrow). Mouth parts include hypostome, chelicera and pedipalps. The pictures have been taken through a ZEISS EVO MA 15, at MEMA (Centro di Servizi di Microscopía Elettronica e Microanalisi, University of Firenze), using the signals produced by secondary electrons, accelerated at 10 KV, with a resolution of 1024x768 nm.

Three biological replicates for each appendage were prepared for ‘reproductive mites’ and for ‘phoretic mites’ from young bees, while a single pool was prepared for ‘phoretic mites’ from foragers or drones, which are more difficult to collect; protein extracted from these latter samples were divided into three aliquots (technical replicates) before enzymatic digestion. The organs were dissected from 35 reproductive and phoretic *Varroa* on young bees, from 50 phoretic *Varroa* on foragers or drones.

### Reagents

All reagents were purchased from Sigma-Aldrich (Milano, Italy) and were of reagent grade. Tris, glycine, Tween-20, urea, nitrocellulose membrane were from Euroclone. Trypsin was from Promega (Sequencing Grade Modified Trypsin) and Lys-C from Thermo Scientific (MS grade). The hand-made desalting/purification STAGE (STop And Go Extraction) tips were prepared using three C18 Empore Extraction Disks (3M)^37,38^.

### Protein extract preparation

Tissues were crushed in a mortar under liquid nitrogen and recovered with 40 μL of 50 mM Tris-Cl buffer pH 7.4, containing 6M Urea/2M thiourea, and centrifuged at 14,000 rpm at 4°C for 30 min. Supernatants were collected and pellets washed with 10 μL of the same buffer, centrifuged at 14,000 rpm at 4°C for 15 min and added to the first supernatants for the analysis. Total protein was measured by the Bradford colorimetric assay, with the “Bio-Rad Protein Assay” kit and Infinite PRO 200 reader (TECAN). Bovine serum albumin was used to generate a standard curve. Protein digestion was carried out on 15 μg protein extracts. Reduction, alkylation and digestion were performed as previously described^39,40^.

The digested samples were then acidified with trifluoracetic acid and desalted on STAGE tips^38^. The eluates were concentrated and reconstituted to 20 μL in 0.5% acetic acid, prior to LC-MS/MS analyses.

### Mass Spectrometric Analysis

Peptide mixtures were analysed on a LC-MS/MS system (Eksigent nanoLC 1D+ coupled to a Q Exactive HF mass spectrometer, Thermo), and 4.5 μg of peptides were injected into a MonoCap C18 HighResolution 2000 (100 micron i.d., 200cm length, GL Sciences). The flow rate was 400nL/min with a gradient from 5% to 60% of solvent B (solvent A= 5% acetonitrile, 0.1% formic acid; solvent B= 80% acetonitrile, 0.1% formic acid) in 255 minutes. The nano-spray source was operated with a spray voltage of 2.1 kV and ion transfer tube temperature of 260 °C. Data were acquired in data dependent mode, with one survey MS scan at the resolution of 60,000 at m/z 400, followed by up to 10 MS/MS on the most intense peaks at 15,000 resolution. Once selected for fragmentation, ions were excluded from further selection for 30 seconds, in order to increase new sequencing events.

### Identification of putative NPC2- and OBP-like Varroa sequences

We used two complementary approaches to detect NPC2- and OBP-like sequences. First, we used local BLAST (v2.6.0; default parameters) to identify *Varroa* protein sequences which are similar to annotated NPC2 proteins in three other arachnids: *Ixodes ricinus* (the castor bean tick), *I. scapularis* (the deer tick) and *A. americanum* (the lone star tick). We repeated this approach using OBP-like sequences that are known in *I. scapularis, A. americanum*, and *Dysdera silvatica.* Next, we retrieved the representative proteome associated with the MD-2-related lipid-recognition (ML) domain (PF02221) – which is found in NPC2 proteins – and used that to query the *Varroa* proteome to find other proteins with this domain. In all cases, we searched for homologs in both the database of known *Varroa* proteins (the same database used by McAfee et al.^41^ to construct the *Varroa* protein atlas) as well as protein sequences generated from a 6-frame translation of the genome sequence (ADDG00000000.2) which were at least 100 residues long. We then merged the results and manually checked the sequences for appropriate length, pattern of cysteines and presence of signal peptide to eventually produce the list of candidates *Varroa* NPC2 and OBP-like sequences. The same inspection was accomplished on sequences reported by Eliash and co-workers^26^ in their transcriptomic analysis of *Varroa* chemosensory tissues. Finally, sequences recognized as OBP-like and NPC2 were checked by running a tblastn against a nucleotide collection (nr/nt) of *V. destructor*.

### Data processing

Raw files of each sample were analyzed using MaxQuant software (version 1.5.8.3)^42^ and the derived peak list was searched with Andromeda search engine^43^. The search was performed against a combined database (available at www.proteomexchange.org; accession: PXD008679) containing 32,122 sequences as follows: annotated *Varroa* protein sequences (kindly provided by Dr. Jay Evans, from the Bee Research Laboratory at the USDA); *Varroa* six-frame translation sequences with supporting peptide data found in McAfee et al.^41^; sequences of all viruses known to infect honey bees and *Varroa*; the few *Varroa* sequences from NCBI published before the latest genome; and the protein sequences of honey bees (OGSv3.2). Since our target was to identify putative soluble olfactory proteins, the final list of our predicted NPC2 and OBP-like sequences were added to the database used for the search (if they were not already present). Default search settings of MaxQuant were used, including trypsin cleavage specificity, 2 allowed missed cleavages, variably oxidized methionine, N-terminal acetylation and fixed carbamidomethyl modification. Parent masses were allowed an initial mass deviation of 4.5 ppm and fragment ions were allowed a mass deviation of 0.5 Da. PSM (Peptide Spectrum Match). Identified protein were filtered using a target-decoy approach at a false discovery rate (FDR) of 1%.

Label-free quantification (LFQ) of proteins was done using the MaxLFQ algorithm integrated into MaxQuant and the ‘match between runs’ option was enabled. For protein quantification, we used the following parameters: 2 as minimum ratio count for “Unique+Razor” peptides (i.e. those exclusively shared by the proteins of the same group), peptides with variable modifications were included, and enabled “discard unmodified counterpart peptide”. The data relative to identification and quantification are contained in the MaxQuant output file named proteinGroups.txt and are reported in Supplementary Table S1.

### Correction of proteomics data for honey bee contamination

*Varroa* is an obligate ectoparasite and feeds on honey bee hemolymph – therefore, its legs and mouthparts are unavoidably contaminated with honey bee proteins. However, the level of contamination is not consistent between samples, presumably because it depends on how recently the mite was last feeding. Using log2 transformed LFQ intensities, we observed that between 6 and 38% of a given sample was composed of honey bee proteins. Since MaxQuant LFQ intensities are scaled against the total ion current, the honey bee contamination will artificially skew the *Varroa* LFQ intensities from highly contaminated samples to be lower than in the absence of contamination. To account for this fact, we applied a correction factor to log2 transformed LFQ intensities of each sample based on its level of contamination (see Supplementary Table S2) prior to differential expression analysis.

### Differential expression analysis

Further analysis of the MaxQuant-processed and corrected data was performed using Perseus software (version 1.5.6.0). First, hits to the reverse database, contaminants and proteins identified only with modified peptides were deleted. LFQ intensity values obtained for the technical replicates of 'phoretic mites’ (from foragers or drones) were averaged and considered as a single biological replicate. Differences in single protein levels were first evaluated between the three appendages, independently from stage, considering only proteins with at least 7 observations (out of 24). Differential expression analysis was performed using ANOVA, where p-values were Benjamini Hochberg corrected at 5% FDR. A post-hoc t-test was applied to determine proteins significantly different between two appendages, using the same correction as in ANOVA.

For differential expression analysis within the same stage, proteins with at least 3 observations (out of 9) for reproductive mites and proteins with at least 4 observations (out of 12) were considered in ANOVA, subjected to Benjamini Hochberg correction at 5% FDR. A post-hoc t-test was then applied to highlight differences between tissues of the same stage. Hierarchical clustering analyses were performed using average Euclideaan distance and the default parameters of Perseus (300 clusters, maximum 10 iterations).

For putative OBP-like and NPC2 proteins, a Wilcoxon signed-rank test with a Monte Carlo simulation was applied, after the imputation (width = 0.3, downshift = 1.8) of missing LFQ values. Differential expression of OBP-like and NPC2 proteins were also evaluated between developmental stages of a previously published *Varroa* proteomics dataset^41^ which included egg, protonymph, female deutonymph, male deutonymph, adult daughter, adult male, and foundress samples (whole mites). The raw data (PXD006072) were searched again with the same MaxQuant parameters as above and the same protein database containing the OBP-like and NPC2 proteins. ANOVA analysis (Benjamini Hochberg-corrected FDR = 5%) was performed in Perseus as above, after filtering for proteins quantified in at least 6 out of 21 processed samples.

### Gene score resampling analysis

We performed a gene score resampling (GSR) analysis to determine if any GO terms were significantly enriched in different tissues and life stages (reproductive and phoretic). We used Blast2GO to retrieve GO terms for *Varroa* proteins using default parameters and the Arthropod protein database for BLAST. We then used the GSR option within ErmineJ (v3.0.3)^44^ with multifunctionality testing enabled, all GO terms (molecular function, biological process, and cellular compartment) included, with the minimum group size set to 3. Enrichment tests were performed using p-values obtained from the differential expression analysis comparing forelegs to second pair of legs as well as mouth parts to second pair of legs (first considering reproductive and phoretic stages together and then considering the two stages separately). Only GO terms that were significant at 10% FDR even after correcting for multifunctionality were considered ‘enriched.’

## Results and Discussion

### Protein expression in tissues

Shotgun proteomic analysis of phoretic and reproductive *Varroa* mouthparts, forelegs, and second pair of legs identified a total of 1189 protein at 1% FDR, of which 231 (about 20%) were honey bee contamination. Since *Varroa* feeds on honey bee hemolymph, such contamination most likely originated from natural interactions between host and parasite, as well as from manipulation during collection. Considering only the *Varroa* proteins, 928, 932, and 908 sequences were identified in the forelegs, second pair of legs and mouth parts respectively. Data regarding the identification of all proteins, together with other information (accessions, scores, percent coverage, missed cleavages, etc.) are reported in Supplementary Table S1. Acquisition methods, databases used, and raw files are available through ProteomeXchange (www.proteomexchange.org; accession: PXD008679).We investigated the number of proteins belonging to different gene ontology (GO categories) to verify whether there were more proteins with chemosensing-related functions in the mouthparts and forelegs (the chemosensory organs) compared to the second pair of legs (a non-chemosensory organ) (Figure 2).

**Figure 2.**
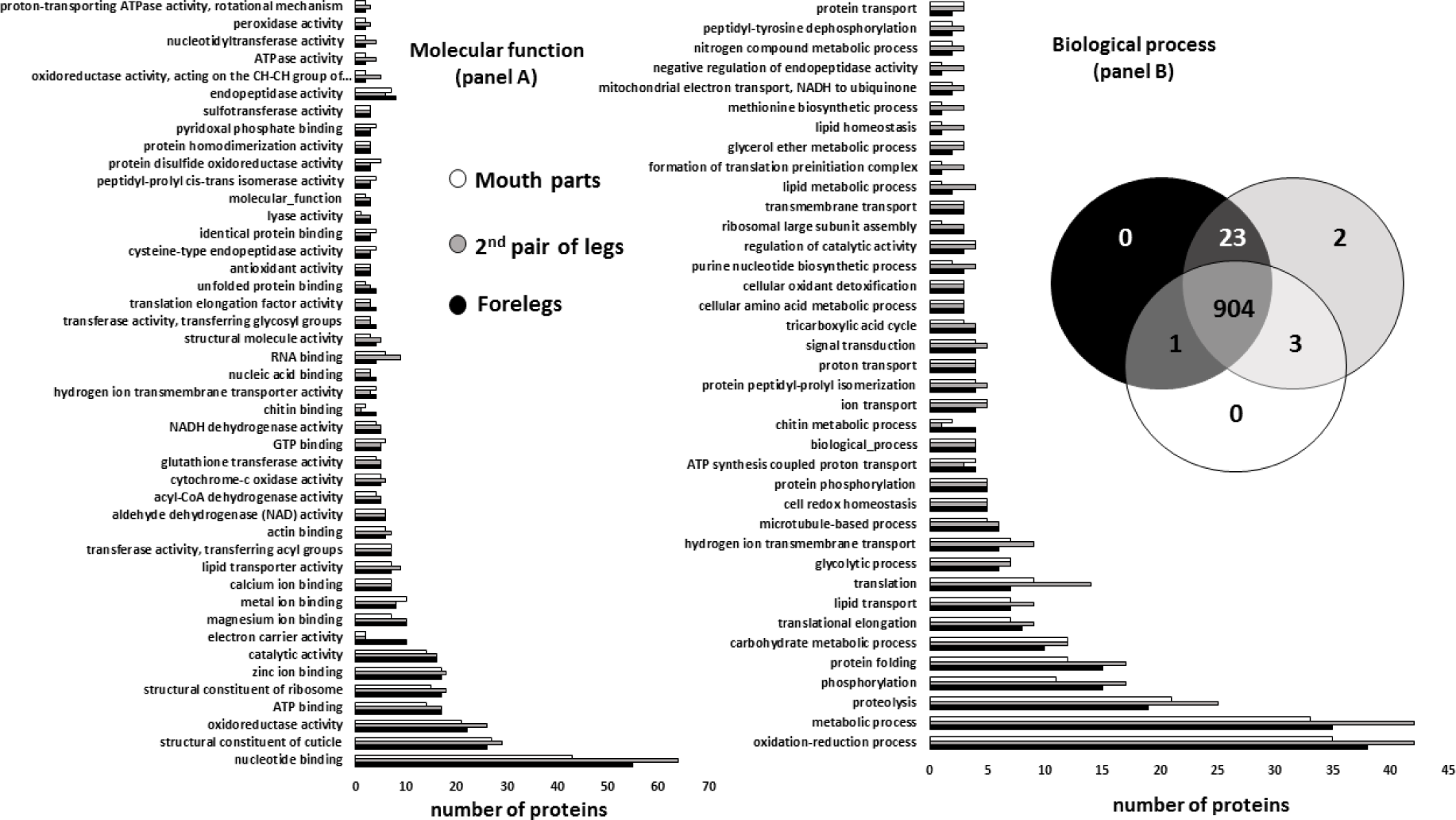
*Varroa destructor* female proteins of forelegs (blue), mouth parts (red) and second pair of legs (green). Panel A: Bar charts reporting GOMF (Gene Ontology Molecular Function) containing at least 4 proteins identified in at least one of the three tissues. Panel B: Bar charts reporting GOBP (Gene Ontology Biological Process) containing at least 3 proteins identified in at least one of the three tissues and Venn diagram based on “Unique+Razor” peptides. In both bar charts we do not observe major differences between the three samples, except for the two molecular function categories ‘electron carrier’ and ‘chitin binding’, that are more represented in forelegs.

The numbers of proteins belonging to each category, both for molecular function and for biological process, are very similar between tissues. Unsurprisingly, the overall most common categories are general GO terms like ‘nucleotide binding’ and ‘oxidation-reduction process,’ while no categories were specific for a particular tissue. In forelegs, two molecular function categories (electron carrier activity and ‘chitin binding) and one biological function (chitin metabolic process) appear to be more represented than in the other tissues, but they are clearly not involved in odorant transport, and no categories were significantly different (Fisher exact test; p value = 0.02). Several proteins with ‘lipid transport’ activity were identified, which includes proteins with hydrophobic binding pockets; however, the numbers were similar between tissues.

The comparable distribution of categories is due to the high degree of overlap in proteins identified in the tissues (Figure 2, panel B). Only two proteins were exclusive to the second pair of legs: an amphiphysin-like isoform X2 and a chaperonin, while no proteins were unique to forelegs or mouth parts. One uncharacterized protein has been found only in the chemosensory tissues, but its function is unknown.

Quantitative differences in protein expression between tissues were evaluated through oneway ANOVA (Benjamini Hochberg-corrected FDR = 5%) followed by a post-hoc t-test. The heatmap reported in Figure 3 shows the 12 proteins differentially expressed among tissues (Table 1).

**Figure 3.**
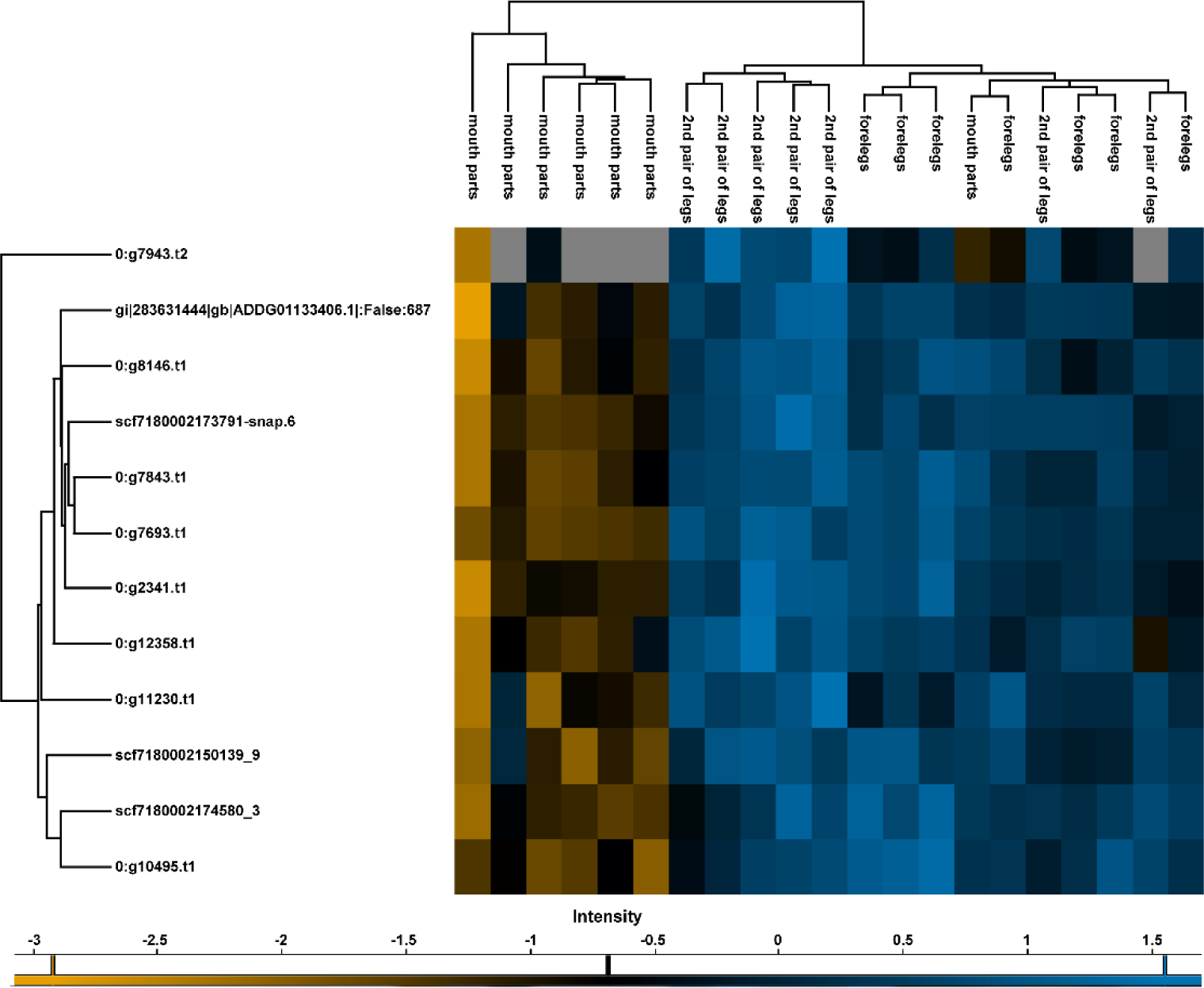
Heatmap representation of the expression of ANOVA significant proteins in the three tissues examined independently from stages. The map has been built making an unsupervised hierarchical clustering (300 clusters, maximum 10 iterations) based on LFQ (Label-free quantification) values of proteins with at least 7 observations resulting significant to ANOVA analysis (Benjamini Hochberg-corrected FDR=5%) among the three tissues. Colour scale reports Z-score log2 transformed LFQ intensity values. Missing data are reported in grey. Major differences are between mouth parts and the other two tissues, that do not show great differences between each other, as displayed in the cluster grouping biological replicates.

**Table 1.** List of proteins differentially expressed among tissues, significant to one-way ANOVA (Benjamini Hochberg-corrected FDR=5%) and to post-hoc t-test.

| Protein IDs<br>(Description) | Peptides | Sequence<br>coverage<br>[%] | Mol.<br>weight<br>[kDa] | -Log<br>ANOVA<br>p value | -Log p-value<br>T-test<br>forelegs_2nd<br>pair of legs | Difference<br>forelegs_2nd<br>pair of legs | -Log p-value<br>T-test mouth<br>parts_2nd<br>pair of legs | Difference<br>mouth<br>parts_2nd<br>pair of legs |
| --- | --- | --- | --- | --- | --- | --- | --- | --- |
| 0:g10495.t1<br>(glutathione S-transferase<br>Mu 3-like) | 13 | 55.8 | 28.4 | 4.37 | 1.06 | 0.42 | 2.80 | -1.26 |
| 0:g11230.t1<br>(voltage-dependent anion-<br>selective channel 2-like) | 14 | 61 | 30.3 | 3.20 | 1.60 | -0.60 | 2.81 | -1.80 |
| 0:g12358.t1<br>(2-oxoglutarate<br>mitochondrial isoform<br>X5) | 17 | 21.6 | 116.6 | 3.00 | 0.38 | -0.25 | 2.46 | -1.42 |
| 0:g2341.t1<br>(pyruvate kinase PKM) | 22 | 53.8 | 56.6 | 3.39 | 0.26 | -0.18 | 2.85 | -1.65 |
| 0:g7693.t1<br>(glycerol-3-phosphate<br>dehydrogenase [NAD(+)]<br>cytoplasmic) | 14 | 53.9 | 34.3 | 4.68 | 0.35 | -0.19 | 3.62 | -1.96 |
| 0:g7843.t1<br>(succinate-- ligase [ADP-forming] subunit<br>mitochondrial-like) | 14 | 36.5 | 47.8 | 3.55 | 0.16 | -0.08 | 2.78 | -1.39 |
| 0:g7943.t2<br>(uncharacterized protein) | 3 | 27.9 | 12.1 | 3.76 | 3.46 | -1.01 | 2.72 | -1.87 |
| 0:g8146.t1<br>(dihydrolipoyl<br>mitochondrial) | 12 | 32.9 | 53.1 | 3.22 | 0.94 | -0.28 | 2.75 | -1.29 |
| gi\|283631444<br>(myosin heavy muscle) | 16 | 62.4 | 26.2 | 3.10 | 0.66 | -0.41 | 2.50 | -2.29 |
| scf7180002150139_9<br>(adenylate kinase<br>isoenzyme 1-like) | 12 | 63.8 | 21.6 | 3.44 | 0.15 | -0.07 | 2.74 | -1.15 |
| scf7180002173791-snap.6<br>(probable citrate synthase<br>mitochondrial) | 15 | 36.4 | 51.8 | 4.29 | 0.72 | -0.29 | 3.29 | -1.77 |
| scf7180002174580_3<br>(secreted salivary gland) | 12 | 57.8 | 23.9 | 4.01 | 0.44 | 0.27 | 2.76 | -1.57 |

Of these, all show significantly lower expression in mouth parts compared to the second pair of legs. Most of the proteins are enzymes, together with one myosin muscle protein, one anion channel, one salivary protein, and one uncharacterized protein. The lower abundance of the salivary protein in the mouthparts is surprising. One explanation could be that the legs might accumulate secreted proteins during the feeding process. Interestingly, four of the differentially expressed enzymes are central to carbohydrate metabolism: pyruvate kinase (0:g2341.t1), glycerol-3-phosphate dehydrogenase (0:g7693.t1), succinate ligase (0:g7843.t1), and citrate synthase (mitochondrial; scf7180002173791-snap.6). Comparing the forelegs and second pair of legs, only one uncharacterised protein showed significantly different expression (0:g7943.t2); this protein does not present conserved domains, nor homologs that could be found in a BLAST search against non-redundant protein sequences of all Arthropoda. None of the significantly different proteins appear to be directly involved in chemosensation.

To determine if total abundances of proteins involved in chemosensation pathways or processes, rather than individual proteins, could be differently represented between tissues, we performed an enrichment analysis by gene score resampling (GSR) based on the t-test p-values (Table 2).

**Table 2.** Go terms significantly enriched in tissues.

|  | Name | ID | # genes | Corrected Pvalue* | Corrected MFPvalue** |
| --- | --- | --- | --- | --- | --- |
| Forelegs vs second pair of legs | phosphorylation | GO:0016310 | 3 | 0.0672 | 0.0693 |
|  | mitochondrion | GO:0005739 | 13 | 0.0709 | 0.0738 |
|  | intracellular membrane-bounded organelle | GO:0043231 | 16 | 0.0826 | 0.0784 |
|  | intracellular organelle part | GO:0044446 | 17 | 0.0666 | 0.0788 |
|  | organelle membrane | GO:0031090 | 12 | 0.0907 | 0.0819 |
|  | protein phosphorylation | GO:0006468 | 3 | 0.0806 | 0.0832 |
|  | phosphorus metabolic process | **GO:0006793** | 7 | 0.0997 | 0.0935 |
|  | phosphate-containing compound metabolic process | **GO:0006796** | 7 | 0.0997 | 0.0935 |
| Mouth parts vs second pair of legs | transferase activity | GO:0016740 | 32 | 0.0262 | 0.0262 |
|  | transferase activity, transferring phosphorus-containing groups | GO:0016772 | 12 | 0.0131 | 0.0568 |
|  | kinase activity | GO:0016301 | 11 | 0.0131 | 0.0721 |

Bold GO terms are redundant with at least one other term
\* p-values corrected for multiple hypothesis testing
\*\* p-values corrected for multiple hypothesis testing and protein multifunctionality

Unlike an over-representation analysis, GSR does not compare a list of ‘significant’ proteins to ‘non-significant’ proteins; rather, the p-values serve as a continuous gene score that can all contribute to the enrichment calculation^43^. Comparing forelegs to the second pair of legs, six significantly enriched GO terms were identified, with phosphorylation (GO:0016310) being the most significantly enriched. Comparing mouthparts to the second pair of legs, three GO terms were significantly enriched, with transferase activity (GO: G0:0016740), particularly phosphorous-containing group transfer activity (GO:0016746) being the most enriched.

### Protein expression within stages

Since different life stages could have different chemosensory needs, we also compared protein expression between the different tissues for reproductive and phoretic mites separately. For example, it is critical for phoretic mites to be able to sense and invade a honey bee cell with a larva at the appropriate age, or else they cannot reproduce; therefore, they could be expressing different proteins to serve this function. We found no differences in protein expression between tissues of reproductive mites (one-way ANOVA, Benjamini Hochberg-corrected FDR = 5%), while 19 proteins were differentially expressed between tissues of phoretic *Varroa* (Figure 4 and Supplementary Table S3). All of those differentially expressed were driven by differences between mouth parts and the second pair of legs. Nine proteins, including three more glycolytic enzymes, are in common with those differentially expressed between tissues, independently from stages.

**Figure 4.**
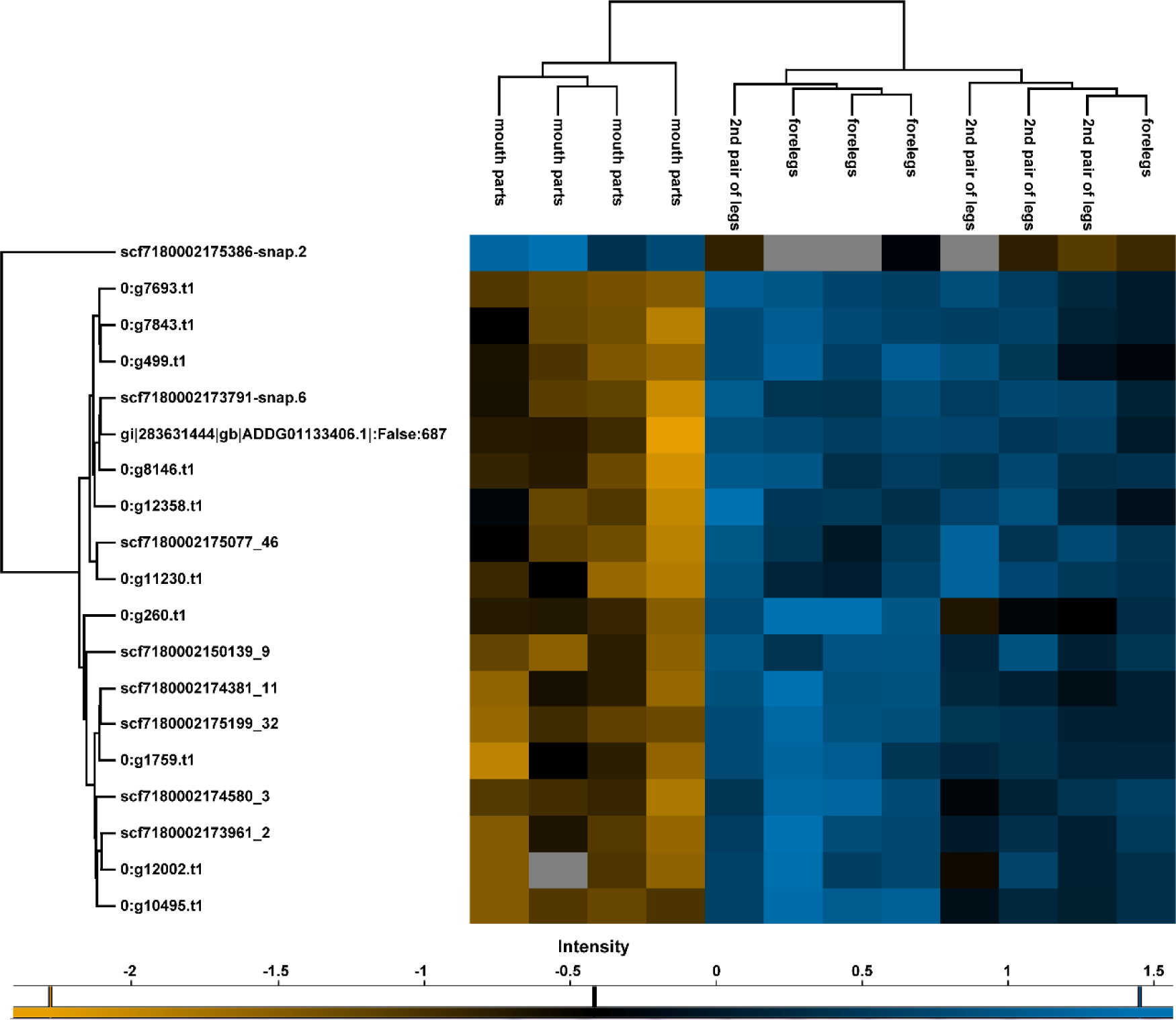
Heatmap representation of the expression of ANOVA significant proteins in the three tissues examined of phoretic mites. The map has been built making an unsupervised hierarchical clustering (300 clusters, maximum 10 iterations) based on LFQ (Label-free quantification) values of proteins with at least 4 observations resulting significant to ANOVA (Benjamini Hochberg-corrected FDR=5%) among the three tissues of phoretic mites. Colour scale reports Z-score log2 transformed LFQ intensity values. Missing data are reported in grey. Major differences are between mouth parts and the other two tissues, that do not show great differences between each other, as displayed in the cluster grouping biological replicates.

To identify cellular processes that may be differently represented between tissues in the two separate stages, we also performed a functional enrichment analysis. Surprisingly, although the phoretic tissue comparison produced more significant expression differences, we found no significantly enriched GO terms between forelegs, second pair of legs or mouth parts. For reproductive mites the “ion binding” category (G0:0043167) was the most enriched in mouth parts compared to forelegs and second pair of legs. These results are reported in Table 3.

**Table 3.**
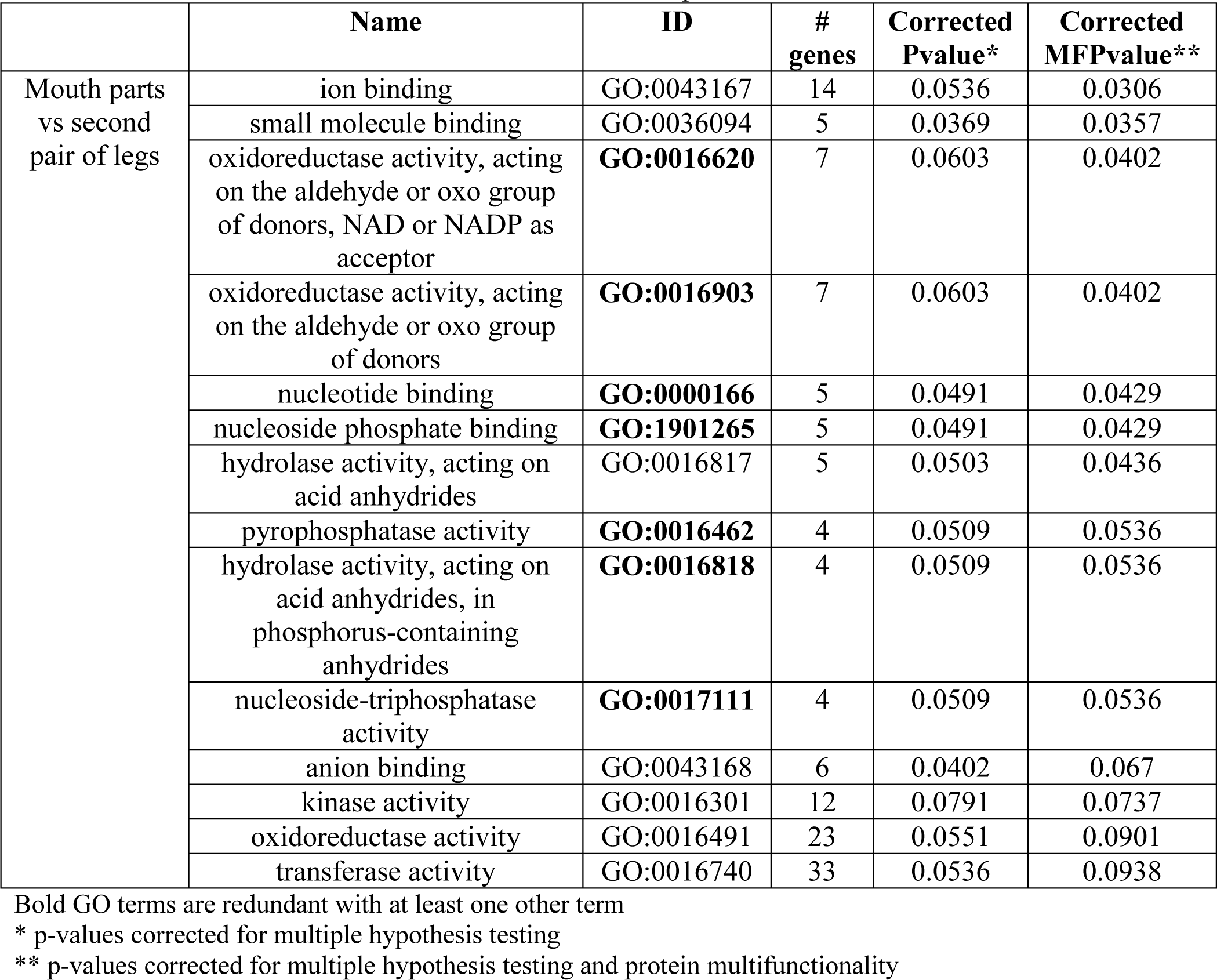
GO terms significantly enriched between tissues of reproductive varroa.

|  | Name | ID | # genes | Corrected Pvalue* | Corrected MFpvalue** |
| --- | --- | --- | --- | --- | --- |
| Mouth parts vs second pair of legs | ion binding | GO:0043167 | 14 | 0.0536 | 0.0306 |
|  | small molecule binding | GO:0036094 | 5 | 0.0369 | 0.0357 |
|  | oxidoreductase activity, acting on the aldehyde or oxo group of donors, NAD or NADP as acceptor | **GO:0016620** | 7 | 0.0603 | 0.0402 |
|  | oxidoreductase activity, acting on the aldehyde or oxo group of donors | **GO:0016903** | 7 | 0.0603 | 0.0402 |
|  | nucleotide binding | **GO:0000166** | 5 | 0.0491 | 0.0429 |
|  | nucleoside phosphate binding | **GO:1901265** | 5 | 0.0491 | 0.0429 |
|  | hydrolase activity, acting on acid anhydrides | GO:0016817 | 5 | 0.0503 | 0.0436 |
|  | pyrophosphatase activity | **GO:0016462** | 4 | 0.0509 | 0.0536 |
|  | hydrolase activity, acting on acid anhydrides, in phosphorus-containing anhydrides | **GO:0016818** | 4 | 0.0509 | 0.0536 |
|  | nucleoside-triphosphatase activity | **GO:0017111** | 4 | 0.0509 | 0.0536 |
|  | anion binding | GO:0043168 | 6 | 0.0402 | 0.067 |
|  | kinase activity | GO:0016301 | 12 | 0.0791 | 0.0737 |
|  | oxidoreductase activity | GO:0016491 | 23 | 0.0551 | 0.0901 |
|  | transferase activity | GO:0016740 | 33 | 0.0536 | 0.0938 |

Bold GO terms are redundant with at least one other term
\* p-values corrected for multiple hypothesis testing
\*\* p-values corrected for multiple hypothesis testing and protein multifunctionality

Overall, surprisingly few proteins were differentially expressed in all our comparisons. The honey bee protein contamination likely interfered with our ability to detect differences in *Varroa* proteins, even with our correction method. In future experiments, more rigorous procedures must be taken to minimize the presence of honey bee proteins (for example, by more efficiently washing the *Varroa* prior to dissection). In addition, proteome depth could likely be improved, which would allow us to detect differences in lower-abundance proteins.

### Putative carriers for semiochemicals

The primary aim of this work was to search for soluble proteins that could represent potential carriers for semiochemicals in *Varroa* and, more generally, in Acari (mites and ticks). Our proteomic analysis on forelegs and mouthparts (which contain chemosensory structures) compared to the second pair of legs (which does not contain chemosensory structures) did not reveal clear differences in proteins, biochemical pathways or processes involved in chemosensation. We therefore chose to use sequence analysis to identify new chemosensory proteins and improve the annotation of those that already exist, then check how these specific proteins were expressed in the different tissues.

OBP-like proteins are a class of soluble proteins identified for the first time in the tick *Amblyomma americanum*^35^ and suggested to be involved in Acari chemodetection. Five transcripts encoding similar proteins have been recently reported in *Varroa*^26^, of which only four can be classified as OBP-like, based on the number and the pattern of cysteines. Using a comprehensive BLAST search strategy (see Methods), we identified two more OBP-like sequences. Figure 5 reports the alignment of the 6 putative OBP-like sequences of *Varroa* together with the two *A. americanum* sequences and one from the tick *I. scapularis*^35^. Although these sequences are very divergent both within and between species, their alignment suggests that they can be all classified in the same family. Sequence XP_022653426.1 of *Varroa*, that appears to be more divergent than the others, likely contains some errors.

The above cited transcriptome work^26^ also reports 8 transcripts proposed to encode NPC2 proteins in *Varroa*. However, after manual inspection, only five of these present the typical pattern of cysteines of NPC2 proteins. In addition, our BLAST search provided one more sequence of the same family. The 6 resulting NPC2 protein sequences of *Varroa* are aligned in Figure 5. Sequences reported in both the OBP-like and NPC2 alignments have been manually corrected for errors at N-term, at C-term or inside the sequences, using Signal-IP 3.0 prediction server, assuming errors at stop codons and/or analysing results from BLAST search between protein and nucleotide sequences (Supplementary file S4). Moreover, no peptide belonging to the above mentioned wrong sequences has been identified in our work.

**Figure 5.**
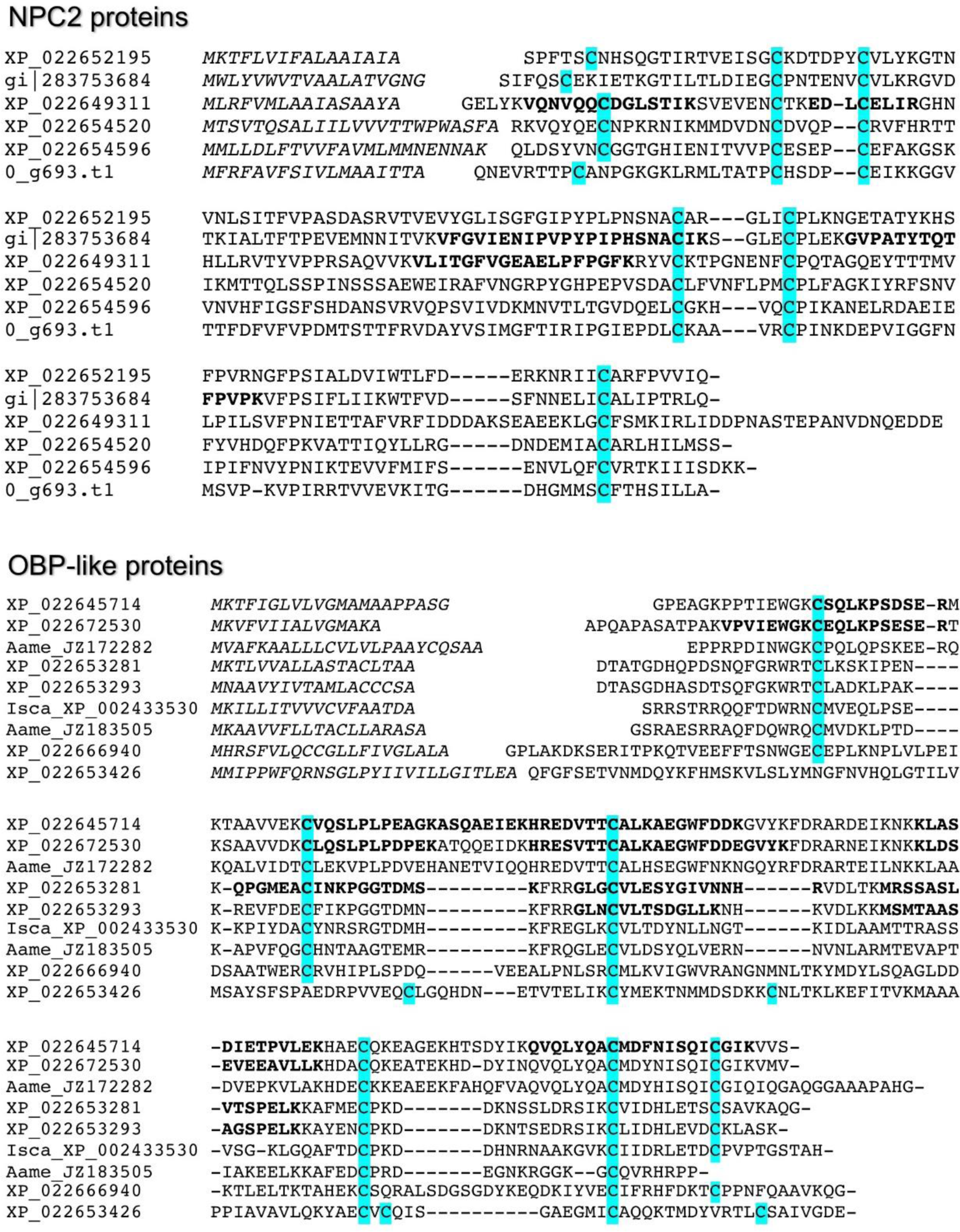
Alignment of protein sequences of *V. destructor* NPC2 (A) and OBP-like (B) proteins. Predicted signal peptides are indicated in italic, while the peptides identified by mass spectrometry are indicated in bold.

Identity values between the six NPC2 sequences of *Varroa* and those of the honey bee never exceed 30% and we only included proteins that could be unequivocally assigned to *Varroa* in the analysis, thus excluding the possibility of contamination. Instead, we found substantial amounts of honey bee OBP13, OBP14 and CSP3. These same proteins had been reported as the only OBPs and CSPs present in honey bee larvae, apart from traces of OBP15^45^; this is consistent with contamination of the *Varroa* sample through larval feeding.

We identified four of the 6 predicted OBP-like proteins in our proteomic analysis, as well as two of the six predicted NPC2 proteins. In order to minimize the effect of possible contamination with bee proteins and assuming that tissue samples dissected out of the same specimens’ pool were contaminated to similar extent, we compared the expression of OBP-like and NPC2 proteins between the three tissues dissected from the same pool by applying a Wilcoxon signed-rank test. The data used for this analysis consists of only these 6 proteins’ log2 transformed LFQ values (with 25% missing values imputed). The abundances of both NPC2 proteins appeared not to change; however, those of two OBP-like proteins, XP_022653293.1 and XP_022653281.1, were higher in forelegs with respect to second pair of legs (respectively z=-2.197, p=0.032; z=-2.028, p=0.046), a result consistent with the forelegs being some of the mite’s main chemosensory appendages. Figure 6 reports, for each protein, the ratio between LFQ values of forelegs and mouth parts with respect to second pair of legs.

**Figure 6.**
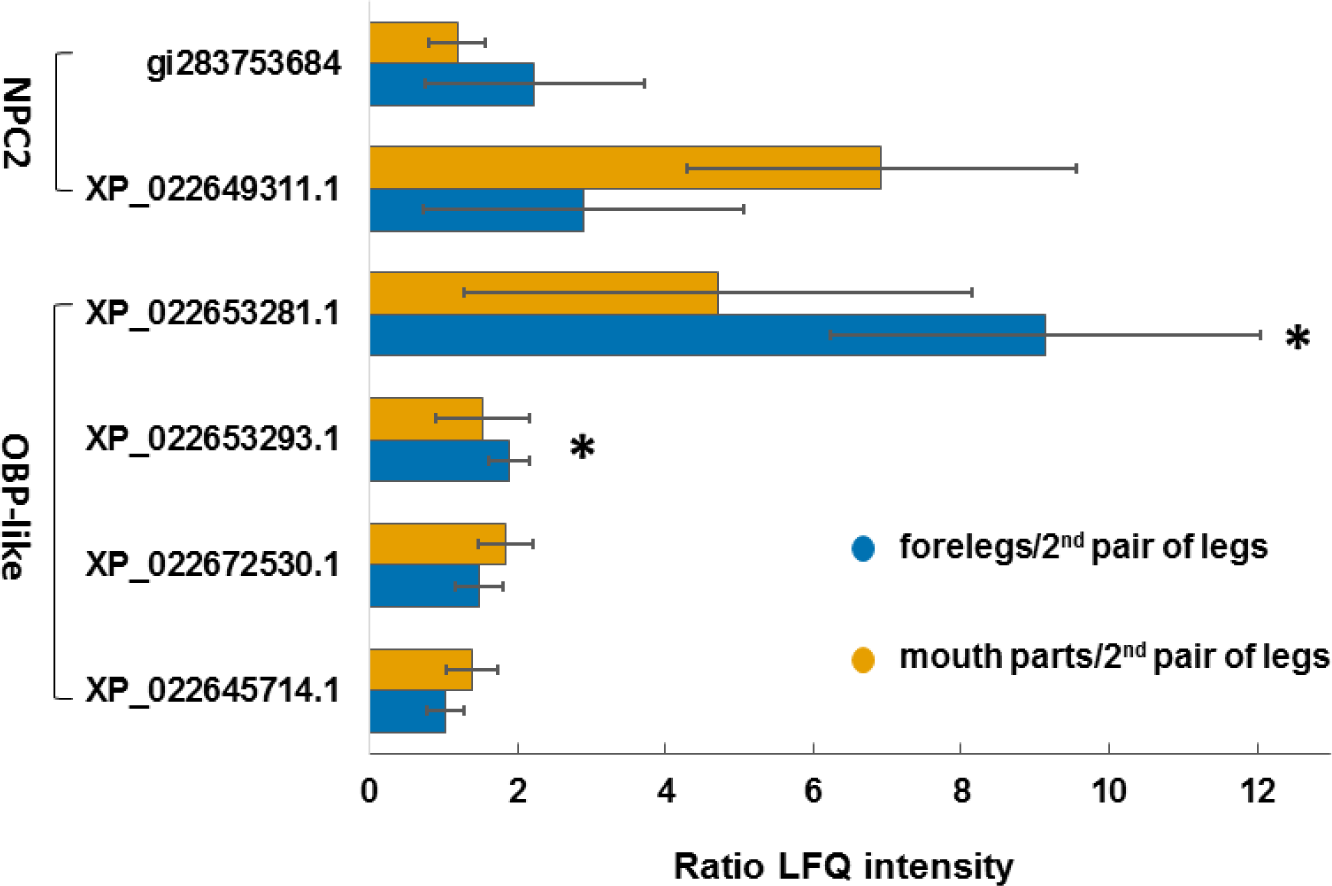
Bar chart reporting the ratio between LFQ intensity values of identified OBP-like and NPC2 of forelegs (blue bars) and mouth parts (orange bars) with respect to second pair of legs. Proteins significant (p<0.05) at Wilcoxon signed-rank test are indicated with an asterisk.

The present proteomics study only analyzes adult female mites; however, chemosensory proteins could also be expressed in males or in other developmental stages. For example, a male mite could require chemosensory abilities to detect when a female is ready for copulation. Therefore, we also evaluated expression of NPC2 and OBP-like proteins identified within the developmental stages of a previously published *Varroa* proteomics dataset^41^. In this analysis, the same OBP-like and NPC2 proteins as reported above were identified, as well as the OBP-like protein, XP_022645714.1. Out of the 5 proteins, two OBP-like were significantly different (XP_022653281.1 and XP_022653293.1; one-way ANOVA; Benjamini Hochberg-corrected FDR = 5%), as reported in Table 4.

**Table 4.**
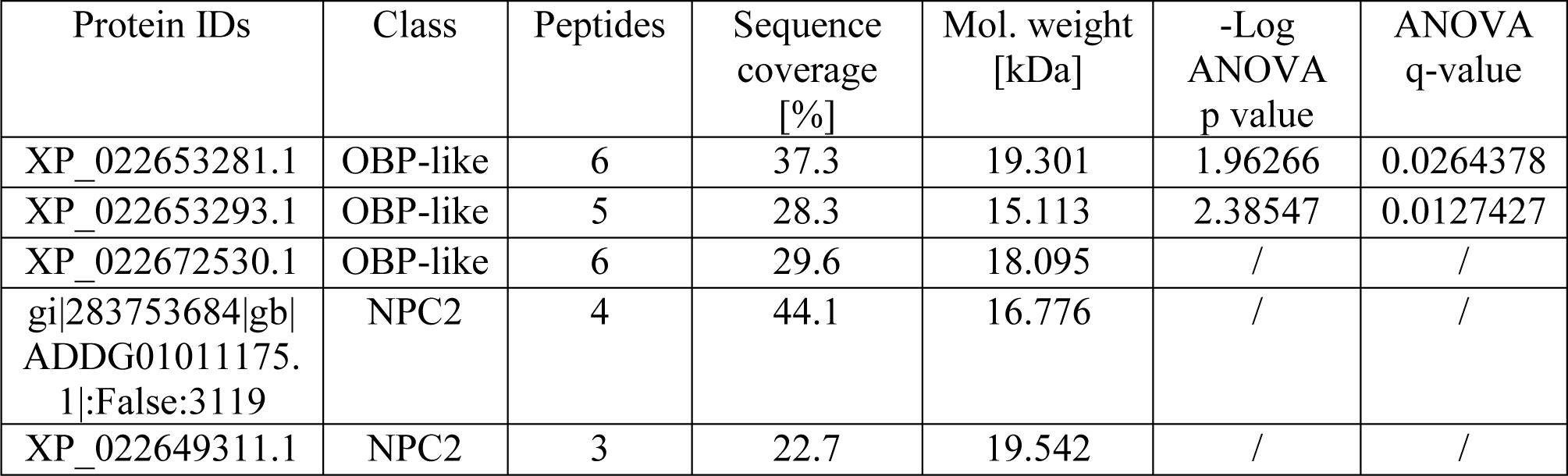
Proteins identified and significant t McAfee et al., 2017).

| Protein IDs | Class | Peptides | Sequence coverage [%] | Mol. weight [kDa] | -Log ANOVA p value | ANOVA q-value |
| --- | --- | --- | --- | --- | --- | --- |
| XP_022653281.1 | OBP-like | 6 | 37.3 | 19.301 | 1.96266 | 0.0264378 |
| XP_022653293.1 | OBP-like | 5 | 28.3 | 15.113 | 2.38547 | 0.0127427 |
| XP_022672530.1 | OBP-like | 6 | 29.6 | 18.095 | / | / |
| gi\|283753684\|gb\|ADDG01011175.1\|:False:3119 | NPC2 | 4 | 44.1 | 16.776 | / | / |
| XP_022649311.1 | NPC2 | 3 | 22.7 | 19.542 | / | / |

Inspecting the abundance of these two proteins, we found that XP_022653281.1 looks egg-biased and expressed only in foundress, within the adult stages, while the protein XP_022653293.1 looks deutonymph/adult-biased. None of the proteins appears to be sex-biased.

## Conclusions

This work presents for the first time a proteomic investigation of chemosensory appendages (forelegs and mouth parts) in *Varroa destructor* adult females at two physiological stages: reproductive and phoretic. The number of identified proteins in these tissues is comparable to the one obtained for the second pair of legs, the control tissue. Differential expression analysis between tissues and within stages revealed several differences in protein expression, but without relation to chemosensing. Moreover, the enrichment analysis by gene score resampling did not show any category clearly involved in odor perception.

An in-depth sequence analysis has allowed us to identify new putative carrier proteins for semiochemicals and an improved annotation of those already reported. In this work we identified protein expression of 4 out of 6 OBP-like sequences, and 2 out of 6 NPC2 sequences of *Varroa.* Unlike what reported for their transcripts expression, at the protein level NPC2 and OBP-like proteins were more abundant in forelegs and mouth parts, bearing the mite’s chemosensory appendages, with respect to second pair of legs. A closer inspection of the abundance of semiochemical carrier proteins, through a paired t-test, revealed that 2 OBP-like proteins were significantly more expressed in forelegs with respect to second pair of legs.

While for NPC2 proteins a function of semiochemical carriers has been supported by ligand–binding experiments and immunocytochemistry, a functional characterization of OBP-like proteins is still needed to clarify their physiological role.

## Acknowledgments

Authors are very grateful to Aldo Baragatti, for kindly providing reproductive *Varroa*, and Federico Cappa and Iacopo Petrocelli, for the help in collecting phoretic mites. We also thank Jay Evans for kindly sharing a *Varroa* protein database. This work was supported by funding from University of Firenze (ex-quota 60%) to FRD; by the 2016 Dott. Giuseppe Guelfi post-doctoral Fellowship from the Accademia Nazionale dei Lincei and by Ente Cassa di Risparmio di Firenze to I. I.

The mass spectrometry proteomics data have been deposited to the ProteomeXchange Consortium via the PRIDE^46^partner repository with the dataset identifier PXD008679.

## Supplementary material

**Supplementary Table S1**. Complete list of proteins identified in proteomic analysis of forelegs, mouth parts and second pair of legs of *Varroa destructor* females. The proteingroups table contains information on the proteins identified in all processed raw-files. Each single row contains the group of proteins that could be reconstructed from a set of peptides.

**Supplementary Table S2**. Correction factors applied to log2 transformed LFQ intensities of each sample.

**Supplementary Table S3**. List of proteins differentially expressed among tissues of phoretic mites, significant to one-way ANOVA (Benjamini Hochberg-corrected FDR=5%) and to post-hoc t-test.

**Supplementary File S4**. OBP-like and NPC2 IDs of peptide sequences predicted from the genome, corresponding accession number in NCBI database and in Eliash and co-workers^26^.

